# Cross-brain transfer of high-performance intracortical speech and handwriting BCIs

**DOI:** 10.64898/2026.01.12.699110

**Authors:** Alisa D. Levin, Donald T. Avansino, Foram B. Kamdar, Nicholas S. Card, Maitreyee Wairagkar, Brandon G. Jacques, Justin J. Jude, Carrina Iacobacci, Bayardo E. Lacayo, Payton H. Bechefsky, Samuel R. Nason-Tomaszewski, Darrel R. Deo, Leigh R. Hochberg, Daniel B. Rubin, Ziv M. Williams, David M. Brandman, Sergey D. Stavisky, Nicholas AuYong, Chethan Pandarinath, Scott W. Linderman, Jaimie M. Henderson, Francis R. Willett

**Author notes:** Co-senior author (Scott W. Linderman, Jaimie M. Henderson, Francis R. Willett).

## Abstract

Intracortical brain-computer interfaces (BCIs) that decode complex movements, such as handwriting and speech, can require substantial training data to achieve high performance. We investigated whether leveraging the neural activity recordings of previous users could reduce this initial data collection burden for new BCI users (an approach we call “cross-brain transfer”). Using intracortical recordings from five BrainGate2 clinical trial participants, we tested cross-brain transfer for both speech and handwriting neural decoders trained and evaluated on general, unconstrained corpora of spoken and written English. We found that cross-brain transfer improved decoding performance when training data from the target user was limited (< 200 sentences), and that dataset-specific input layers to the decoder were critical for combining data across users. Without trainable input layers, transfer failed and performed worse than training from scratch on target user data only. Finally, we measured the effectiveness of cross-brain transfer relative to training with (1) more data from the same user and (2) more electrode-permuted data from the same user, which simulates sampling from another brain with identical neural latent structure. In some cases (T16 speech, T12 handwriting), cross-brain transfer appeared as effective as additional permuted data from the same user, while in others (T12 speech, T15 speech) electrode-permuted data was more beneficial. Our results successfully demonstrate and characterize cross-brain transfer learning between multiple intracortical BCI users, for both speech and handwriting, using a general open-ended dataset not restricted to small sets of words or phrases. This work highlights a promising path towards addressing a key barrier to the clinical translation of BCIs, while clarifying when cross-brain transfer may be most beneficial and the decoder design choices needed to realize those gains.

## Introduction

Recent demonstrations have shown that brain-computer interfaces (BCI) can restore high-speed, accurate communication by translating neural signals during attempted handwriting, speech or typing into text^1–6^. However, these brain-to-text BCIs have sometimes required new users to engage in extensive supervised data collection, taking as much as 10 days of data to attain near-peak decoding performance. One way to decrease training time could be to pre-train neural decoders on data from other users and then fine-tune the decoder for a new user, a technique we call “cross-brain transfer.” Transferring models between users could reduce the amount of data needed to adapt to a new user—assuming that users share at least some underlying structure in their neural activity patterns. Although some studies have reported good decoding accuracies with only a small number of training sentences from a single user^5,6^, successful cross-brain transfer would enable more consistently short training data collection times across all users.

In recent years, several studies have reported promising transfer learning results on neural data from non-human primates between different time points as well as between different subjects and/or tasks^7–9^. In humans, a handwriting decoder trained on surface electromyography (sEMG) signals from over 6,500 users of 16 bipolar-channel sEMG bands could generalize to new users without any user-specific training (median character error rate of 4.8% across 50 heldout users)^10,11^, and a typing decoder trained on sEMG signals of 100 sEMG band users could be successfully transferred to others via fine-tuning (mean character error rate of 6.95% across 8 heldout users)^10,11^. More recently, Spalding et al., 2025 demonstrated a cross-subject transfer method for high-density micro-electrocorticography (μECoG) recordings during speech using canonical correlation analysis (CCA) to align neural latent spaces (i.e., the lower-dimensional representations of neural activity) across individuals using a small vocabulary of 52 words, increasing mean phoneme decoding accuracy between 9 phonemes from 24% (patient-specific baseline) to 31% (cross-patient aligned)^12^. Singh et al., 2025 also demonstrated a cross-subject transfer of stereo-electroencephalography (sEEG)-based speech decoders using recurrent neural networks (RNNs) on a small set of 64 possible phrases, decreasing the median phoneme error rate from 57% (within-subject baseline) to 45%, when fine-tuning a model trained on four other subjects^13^.

Despite these advances, no prior work has explored transfer learning between human BCI users during complex motor behaviors, such as handwriting or speech, using intracortical recordings. Moreover, demonstrations of human BCI transfer learning with other neural recording modalities have been limited to small vocabularies rather than open-ended phoneme or character decoding. Unlike classical 2D cursor-control tasks typically controlled with attempted joystick movements^14,15^, these more fine-motor behaviors have historically required substantially larger datasets to saturate decoding performance (e.g., 10 hours^3^ vs. 2 minutes^16^). Here, we evaluate the benefits and limitations of cross-brain transfer using a large dataset (48.9 hours, 5 participants) of attempted handwriting and speech movements from open-ended corpora of spoken and written English, thus tackling this problem in the more clinically-relevant general vocabulary setting for the first time.

## Results

### Cross-brain transfer improves decoding accuracy when training data is limited

To investigate the potential benefits of transfer learning between intracortical BCI users during complex motor behaviors, we leveraged a library of neural recordings collected while participants performed speech or handwriting instructed delay tasks (Fig. 1a). This dataset spans five BrainGate2 clinical trial participants: T5, T12, T15, T16 and T18, each of whom had microelectrode arrays placed in speech- or upper extremity-related areas of the precentral gyrus (Fig. 1b–c). With these neural recordings, we first tested whether including data from additional source users as decoder training data (“cross-brain transfer”) improved performance relative to training on a single target user’s data alone. We evaluated decoder performance offline as a function of the number of target user sentences included as training data, in order to assess whether cross-brain transfer was more helpful in a data-limited regime (Fig. 2). We found that cross-brain transfer improved performance when training data was limited (< 200 sentences). This result was consistent across decoders for all speech and handwriting target users, except for participant T18. Cross-brain transfer may not have been useful for T18 since performance on decoders trained from scratch already appeared to saturate with only ∼20 sentences of T18 training data, possibly due to T18’s unusually high signal quality.

**Fig. 1.**
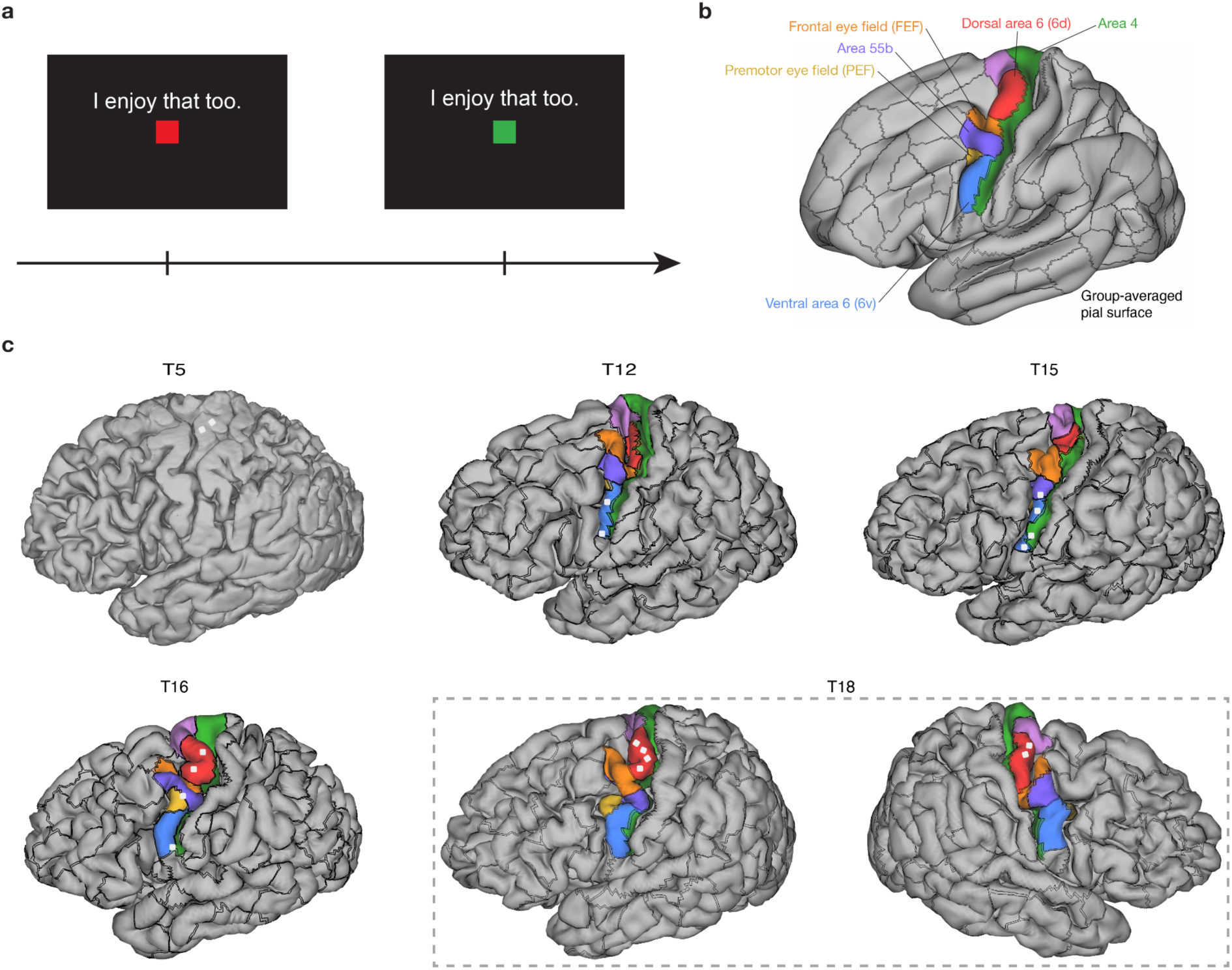
Speech and handwriting task design and electrode array placements for each participant. **a,** Neural recordings were collected from participants during speech or handwriting instructed delay tasks. **b,** Human Connectome Project (HCP) brain regions within the precentral gyrus are displayed on the group-averaged atlas brain (pial surface) for reference. **c,** Human Connectome Project (HCP) cortical parcellations for participants T12, T15, T16, and T18. Each participant’s pial surface is pictured with areal borders (black lines) of the parcellated regions, with regions of interest along the precentral gyrus shown in color. Brain region borders were estimated using presurgical structural and functional (resting state) MRI data. For participant T5, whose arrays were placed without conducting the HCP protocol, a pial surface without parcellated regions is pictured. The white squares indicate array placement locations, which were estimated using photographs taken during surgery. See Willett et al., 2021 for details on participant T5, Deo et al., 2024 for details on participants T12, T15 and T16, and Jude et al., 2025 for details on participant T18.

**Fig. 2.**
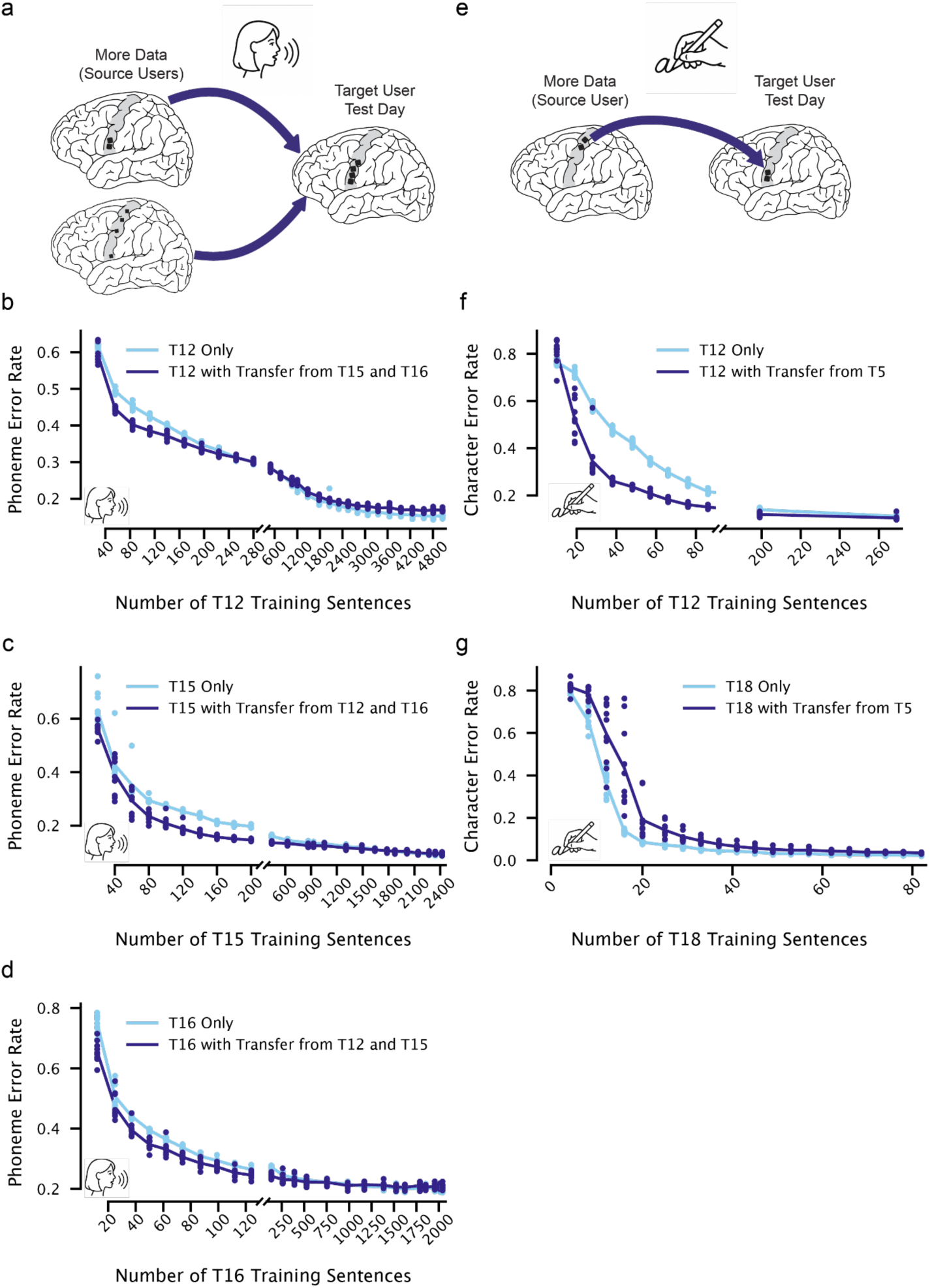
Leveraging other users’ neural data improves decoding accuracy for new users when new training data is limited. **a,** Diagram of cross-brain transfer approach for speech. A large library of source data from two users was used to train a base decoder, which was then fine-tuned on data from a single test day from a new target user. **b,** Speech decoding performance as a function of the number of training sentences from the target user, participant T12. Lines show the mean phoneme error rate across decoder seeds, and points show individual decoder phoneme error rates. Decoders trained from scratch on T12’s data are shown in light blue. Decoders first trained on speech sessions from participants T15 and T16 and then fine-tuned on the T12 data are shown in indigo. Points to the left of the x-axis break include training data only from T12’s test day, while points to the right of it include data from additional T12 days. Decoder performance was always evaluated on the same held-out test set from T12’s test day. **c,d** The same as **b**, but for T15 and T16 as the target users. **e,** Same as **a,** but for handwriting and with only one source user. **f,g** The same as **b–d**, but for handwriting.

### Fitting unique, dataset-specific input layers was necessary for successful cross-brain transfer

Our RNN decoding architecture used trainable, session-specific input layers to more effectively aggregate data from different users and from different days within the same user. We investigated whether these input layers were necessary for successful cross-brain transfer by comparing performance with and without session-specific input layers, in a data-limited regime where cross-brain transfer was shown to be helpful. We found that when RNNs were trained with a single shared, learnable layer across all users and days, performance was often much worse than even training on the target user’s data alone (Fig. 3). Thus, the learnable input layers not only improved performance in the presence of neural non-stationarities across sessions within a single user, as demonstrated in prior work^1,3,17^, but also aided in combining neural data across different users. This suggests that specific mechanisms like unique, session-specific layers may be necessary for cross-brain transfer to improve performance with intracortical neural data.

**Fig. 3.**
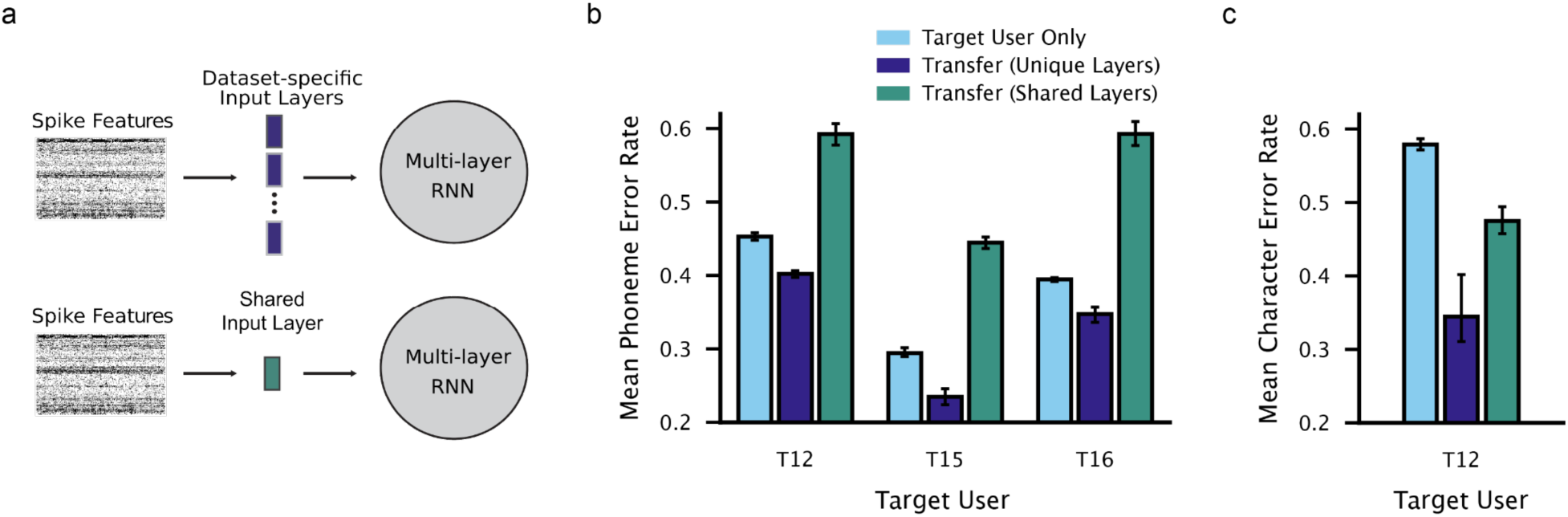
Fitting unique, session-specific input layers was necessary for successful transfer. **a,** Diagram highlighting our standard decoder architecture with unique, session-specific input layers, contrasted with an ablation to that decoder which instead uses a single, shared learnable layer for all data sessions across all users. **b,** A comparison of speech cross-brain transfer performance with and without session-specific input layers (indigo and teal bars, respectively). Bar heights indicate the mean phoneme error rate across decoder seeds (error bars denote 95% confidence intervals). A small number of training sentences were used, with the number chosen based on results from Fig. 2 (T12: 84, T15: 80, T16: 50). The performance of decoders trained on the same data from scratch is also shown for reference (light blue bars). **c,** Same as **b,** but for handwriting and with T12 as the target user (28 sentences were used for training data).

### Effectiveness of cross-brain transfer compared to electrode-permuted data

We next examined the effectiveness of cross-brain transfer relative to two controls: training with more data from the same user, and training with more electrode-permuted data from the same user. The electrode permutation control assesses whether leveraging data from other source users is comparable to using data that shares identical neural latent structure with the target user (i.e., differs only by an orthonormal transform). As a baseline, we evaluated the performance of a decoder trained from scratch on only a single test day of target user data (light blue condition). We then compared this to a decoder initially trained on more target user days (purple) and then fine-tuned on the target user’s test day. Finally, we also compared a decoder pre-trained on electrode-permuted versions of those target user days (purple with hash) as well as to cross-brain transfer (indigo). For some users (T16 speech, T12 handwriting), cross-brain transfer was as effective as additional permuted data from the same user (Fig. 4d–e). For other users (T12 speech, T15 speech), electrode-permuted data was more beneficial than cross-brain transfer (Fig. 4b–c). The results for T12 and T15 suggest that, at least for some users, their recorded neural activity patterns may substantially differ from an orthonormal transform of other users.

**Fig. 4.**
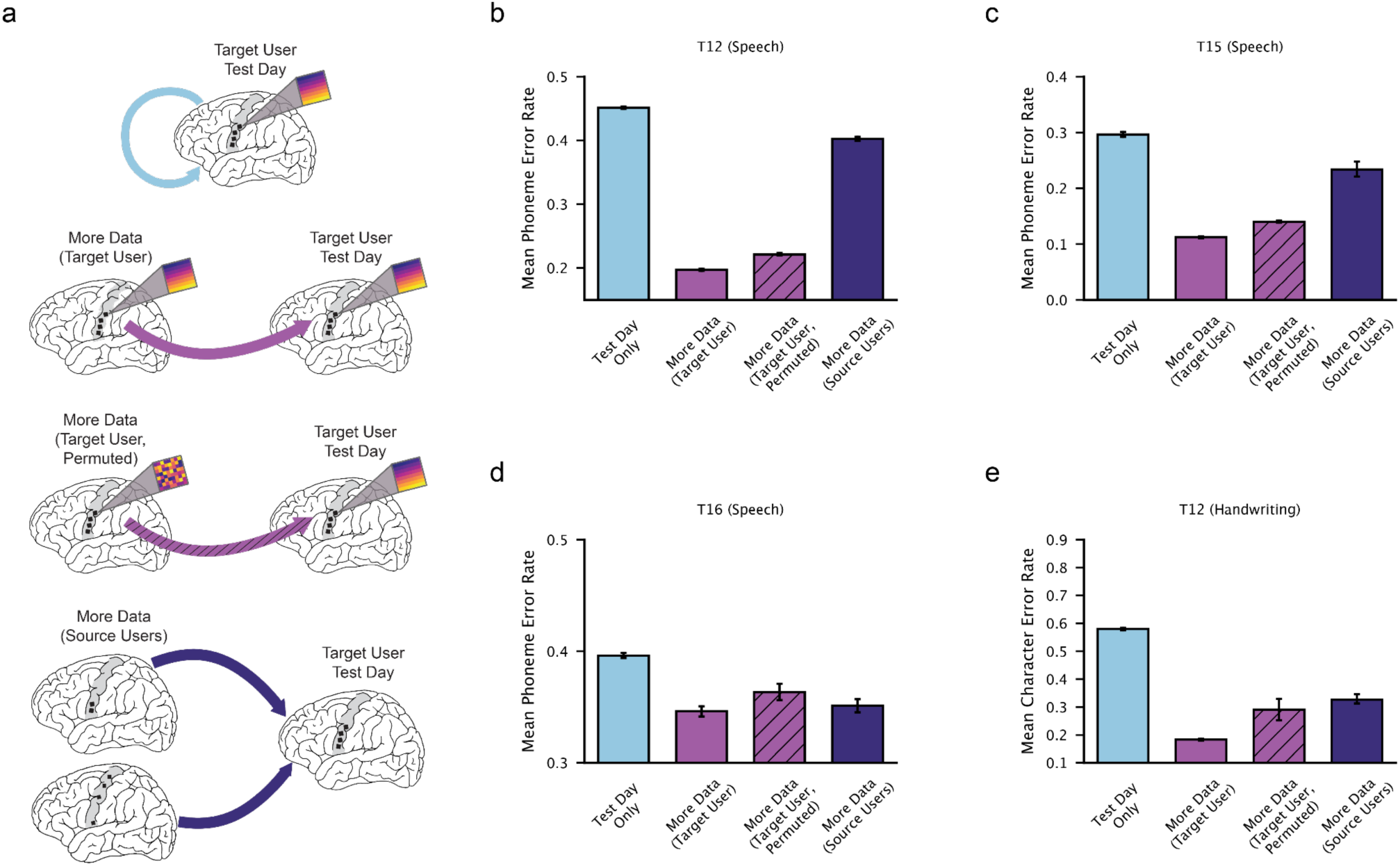
Effectiveness of cross-brain transfer compared to electrode-permuted data. **a,** Diagrams illustrating transfer to the target user’s test day using (top) more data from that target user, (middle) permuted data from that target user, and (bottom) data from other source user(s). **b,** A comparison of T12 speech decoding performance when: training from scratch on 84 sentences of training data only (“test day only”); using a base decoder initialized on additional T12 data (“more data”) and fine-tuned on the same 84 sentences; using a fine-tuned base decoder initialized on additional electrode-permuted T12 data (“more data, permuted”); and using a fine-tuned base decoder initialized using data from the other two source users (“more data, other users”). Bar heights show mean phoneme error rates across seeds (error bars denote 95% confidence intervals). **c,** Same as **b**, but with T15 as the target user (80 training sentences). **d,** Same as **b**, but with T16 as the target user (50 training sentences). **e,** Same as **b**, but for handwriting and with T12 as the target user (28 training sentences).

## Discussion

In this work, we demonstrated the first successful transfer of intracortical speech and handwriting decoders between users. We also showed that the gains from transfer depend on decoder architecture—in our case, the learnable session-specific input layers. Without trainable input layers, training on multiple users not only failed to improve performance, but often made performance worse. We found that cross-brain transfer improved decoding performance when training data was limited (< 200 sentences) for all target users except participant T18 (Fig. 2). Interestingly, we achieved a 2% character error rate for T18 with a decoder trained from scratch on only a single handwriting data collection session. Thus, it is possible that the neural signal was so informative for handwriting decoding that a small amount of T18’s data was sufficient to saturate decoding performance, and cross-brain transfer was therefore unhelpful even in a data-limited regime.

The decoding performance improvements that we observed from cross-brain transfer were modest, consistent with results reported in other recent transfer learning work across various neural recording modalities and algorithmic approaches^7–9,12,13,18^. The studies most closely related to ours are Spalding et al., 2025 and Singh et al., 2025, which demonstrated successful cross-subject transfer for speech using μECoG and sEEG, respectively. Our work extends these studies by using a more general dataset consisting of 48.9 hours of general English as compared to 52 words (Spalding et al.) or 64 phrases (Singh et al.). Decoding performance on small vocabularies cannot be expected to generalize to larger vocabularies, as prior work has shown that a small vocabulary of 50 words is much easier to decode than a large 125,000 word vocabulary^3,5^. As compared to Spalding, we also use a general decoding framework capable of decoding any of 39 phonemes as building blocks for words using a deep learning model, and trainable input layers that do not require matching conditions for CCA-based alignment.

We hypothesize that a key limiting factor to performance gains from cross-brain transfer is not only the amount of training data that one has per user, but—perhaps more importantly—the number of different users whose data can be combined to train generalizable decoders. Meta’s Reality Labs, for example, recently demonstrated the zero-shot generalization of a 16 bipolar-channel surface electromyography (sEMG) decoder by training on data from over 6,500 users^10,11^. This large-scale library of users allowed Reality Labs to both capture inter-individual variability, and train a large neural network better suited to transfer than the smaller data-scarse decoders commonly used for BCIs. Moreover, sEMG time series are comparatively low-dimensional and likely more stereotyped across able-bodied individuals than intracortical signals recorded by hundreds of microelectrodes placed in different cortical regions and clinical populations (amyotrophic lateral sclerosis (ALS), brainstem stroke, cervical spinal cord injury).

Consistent with the hypothesis that data from a large number of users may be needed for more effective cross-brain transfer, we found that, for some target users (T12 speech, T15 speech), training on data from other users was much less effective at improving decoder performance than training on additional, electrode-permuted data from the same user (Fig. 4). Since the electrode-permuted data serves as a simulation of sampling from another brain with identical neural latent structure to the target user, this result implies that neural recordings from these different users are not simply different projections of the same underlying neural representations, and that the main roadblock to cross-user transfer in this case may be differences in the underlying neural latent structure across users. To improve cross-brain transfer, a large library of data from many users may be needed such that for each new user, there is at least one user in the library with similar neural latent structure. While recent studies have focused on methods for linear alignment of cross-user datasets^19–21^, our results suggest that underlying differences in neural encoding and dynamics across users may limit the benefit and applicability of linear re-alignment.

In conclusion, this work demonstrates that cross-brain transfer is a promising approach for reducing the data collection burden for new BCI users. In addition to collecting data from more users, further improvements could be made by exploring other decoding architectures and training techniques, including transfer learning methods such as meta learning^22,23^ or low-rank adaptation^24^, as well as other neural network architectures such as transformers. Although we have found that for smaller datasets, it is difficult to outperform our RNN architecture^3^, larger datasets may enable more elaborate architectures to improve cross-brain generalization and performance in the future.

## Methods

### Participants and Approvals

This study includes data from five participants (referred to as T5, T12, T15, T16 and T18), all of whom gave informed consent and were enrolled in the BrainGate2 Neural Interface System clinical trial (ClinicalTrials.gov Identifier: NCT00912041, registered June 3, 2009). This pilot clinical trial was approved under an Investigational Device Exemption (IDE) by the US Food and Drug Administration (Investigational Device Exemption #G090003). Permission was also granted by the Institutional Review Boards of Stanford University (protocol #52060), University of California, Davis (protocol #1843264), Emory University (protocol #STUDY00003070), Boards of Massachusetts General Hospital (protocol #2009P000505), Brown University (protocol #0809992560) and the VA Providence Healthcare System (IRB-2011-009). All research was conducted in accordance with relevant guidelines and regulations.

T5, a right-handed man with a cervical spinal cord injury (C4 AIS-C), had two 96-channel microelectrode arrays (NeuroPort, Blackrock Microsystems, Salt Lake City, UT) placed in the hand knob^25^ area of the left hemisphere; see Willett et al., 2021 for details. T5 was 69 years old at the time of data collection, and the data reported are from post-implant days 2100–2135.

T12, a right-handed woman with slowly-progressive bulbar-onset Amyotrophic Lateral Sclerosis (ALS), had four 64-channel microelectrode arrays placed in her left hemisphere: two in HCP-identified^26^ area 6v of ventral precentral gyrus (orofacial motor cortex) and two in area 44 of inferior frontal gyrus (considered to be part of Broca’s area); see Willett et al., 2023 for details. T12 was 67–68 years old at the time of data collection, and the data reported are from post-implant days 30–573.

T15, a left-handed man with ALS, had four 64-channel microelectrode arrays placed in his left hemisphere: two in HCP-identified area 6v of ventral precentral gyrus (orofacial motor cortex), one in area 55b, and one in area 4 (primary motor cortex); see Card et al., 2024 for details. T15 was 45 years old at the time of data collection, and the data reported are from post-implant days 27–97.

T16, a right-handed woman with tetraplegia and dysarthria due to a pontine stroke, had four 64-channel microelectrode arrays placed in her left hemisphere: two in HCP-identified area 6d (hand knob), one in area 6v of ventral precentral gyrus (orofacial motor cortex), and one on the border of the premotor eye fields (PEF) and speech-related 55b; see Kunz et al., 2025 for details. T16 was 52 years old at the time of data collection, and the data reported are from post-implant days 32–284.

T18, a right-handed man with a cervical spinal cord injury (C4 AIS-C), had six 64-channel microelectrode arrays placed: four in HCP-identified left area 6d (hand knob) and two in right area 6d; see Jude et al., 2025 for details. T18 was 48 years old at the time of data collection, and the data reported is from post-implant day 162.

### Functional MRI speech lateralization and array placement targeting

Prior to surgery, participant T5’s hand knob area was identified by pre-operative magnetic resonance imaging (MRI)^27^. Participants T12, T15 and T16 underwent pre-operative anatomic and functional brain imaging for speech and language lateralization, surgical planning and array placement targeting (see Willett et al., 2023, Card et al., 2024, and Deo et al., 2024 for array location estimates and further details). Taking advantage of the imaging methods used for participants with arrays in speech-related areas, participant T18 also underwent anatomic and functional brain imaging for surgical planning and hand knob area array placement targeting^6^. See Fig. 1c for the array locations of each participant.

### Neural signal processing

Neural signals were recorded from the microelectrode arrays using the Neuroplex-E system (Blackrock Microsystems) and transmitted via a cable attached to a percutaneous connector. Signals were analog filtered (4th order Butterworth with corners at 0.3 Hz to 7.5 kHz), digitized at 30 kHz (250 nV resolution), and fed into a custom MATLAB Simulink or Python-based BRAND software^28^ for digital filtering and feature extraction. For participants T5 and T12, as well as for participant T16 on session 2024.04.19 and onwards (see Table I for a list of all T16 sessions), digital filtering began with a highpass filter (250 Hz cutoff) that was applied non-causally to each electrode, using a 4 ms delay, in order to improve spike detection^29^. For participants T15 and T18, as well as for participant T16 during all sessions up through session 2024.03.11 (see Table I), digital filtering instead began with a bandpass filter (corners at 250Hz and 5kHz) that was applied non-causally to each electrode, using a 1 ms delay. For all participants, linear regression referencing (LRR)^30^ was then applied to each array-group of 64 electrodes in order to reduce noise artifacts. Then, for all participants, two estimates of neural ensemble activity were computed for each electrode in either 10 ms or 20ms bins. Binned spike band power was computed by taking the sum of squared voltages at each time bin, and threshold crossing rate features were computed by counting the number of times that the filtered voltage time series crossed an amplitude threshold set at either -4.5 (T5, T12, T15, T16) or -3.5 (T18) times the standard deviation of the voltage signal. Threshold crossing rates and spike band power features measure local spiking activity and have been shown to yield similar decoding accuracies and neural population structure as sorted single neurons^31–34^.

**Table I.**
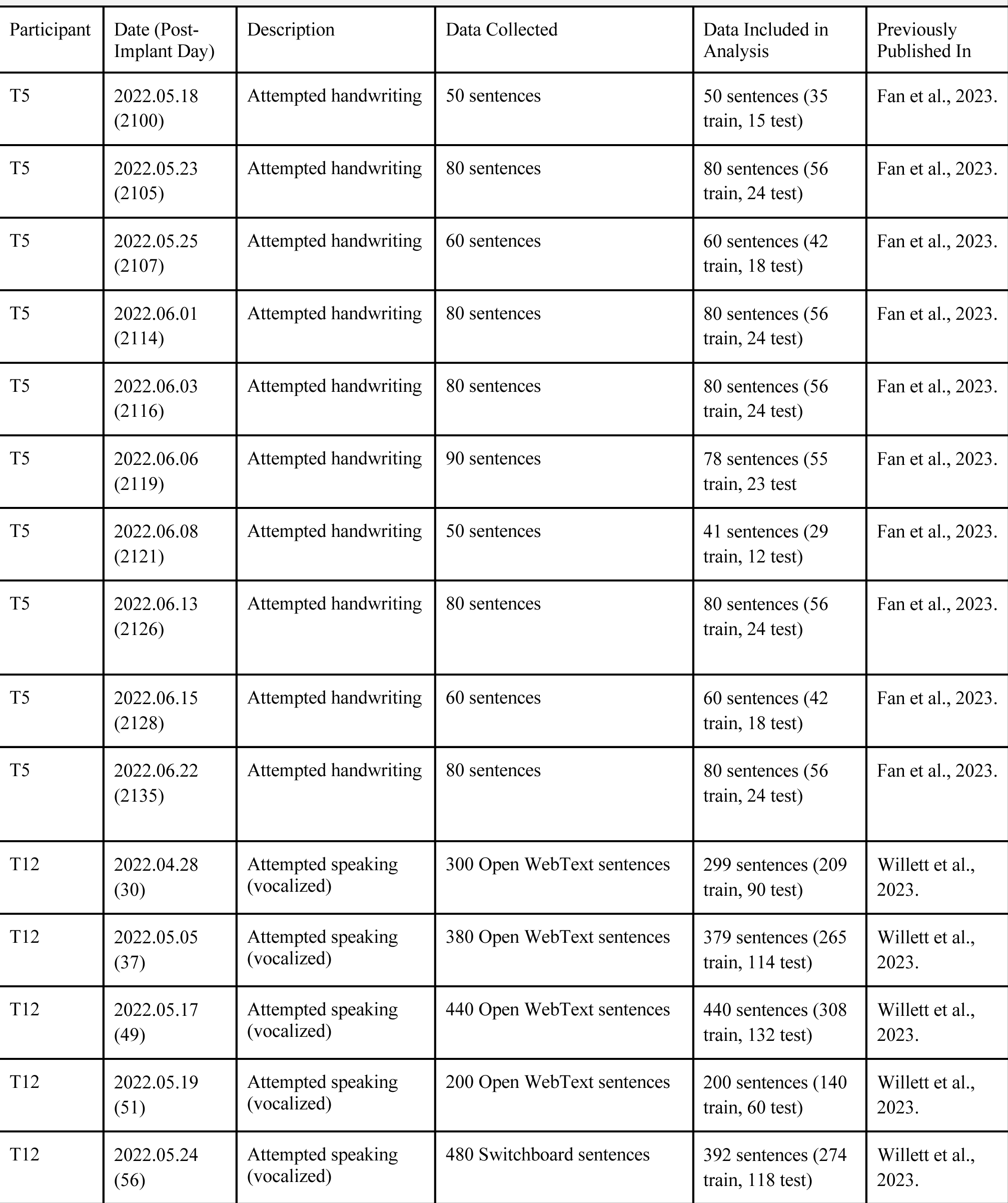
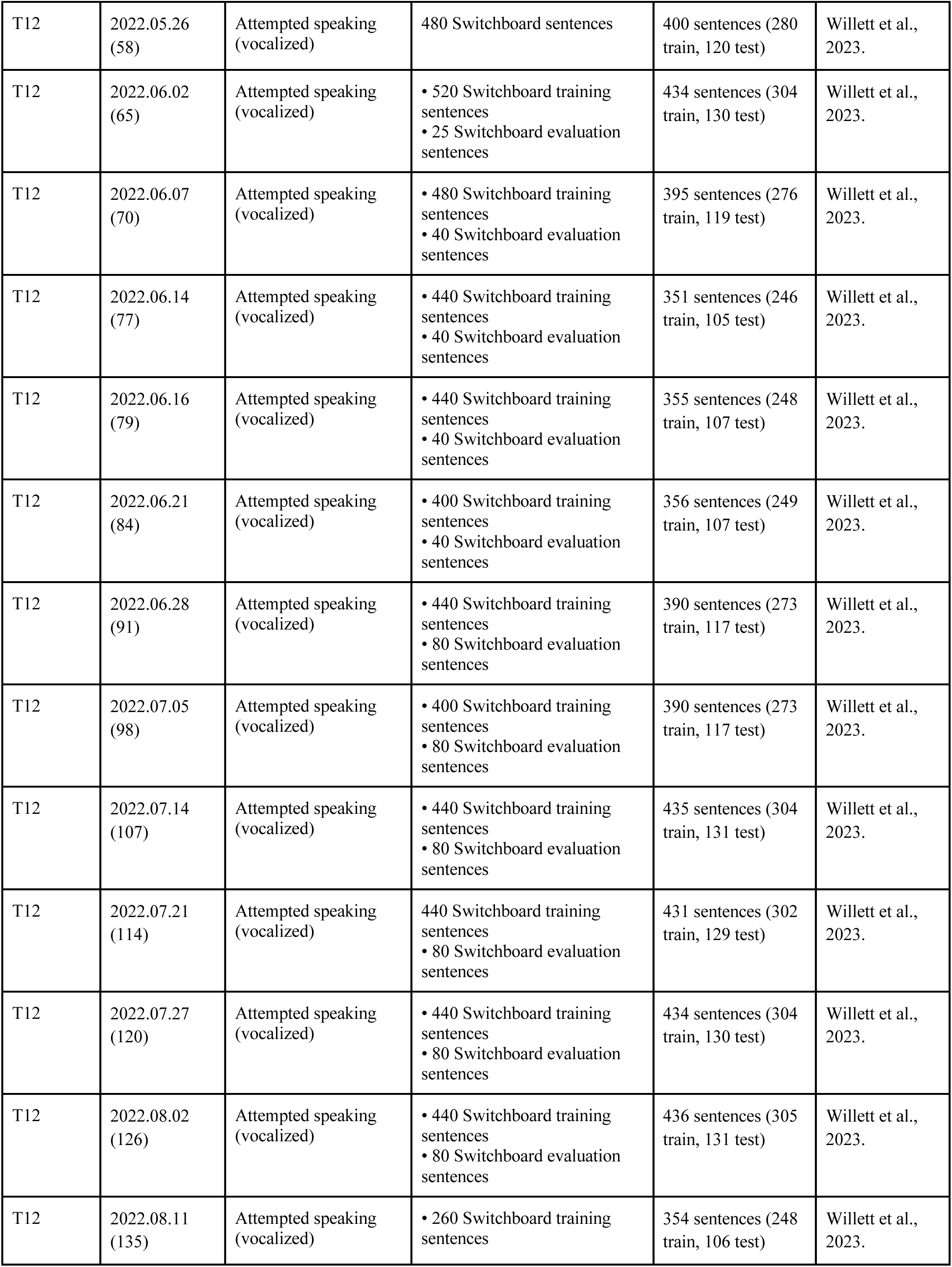
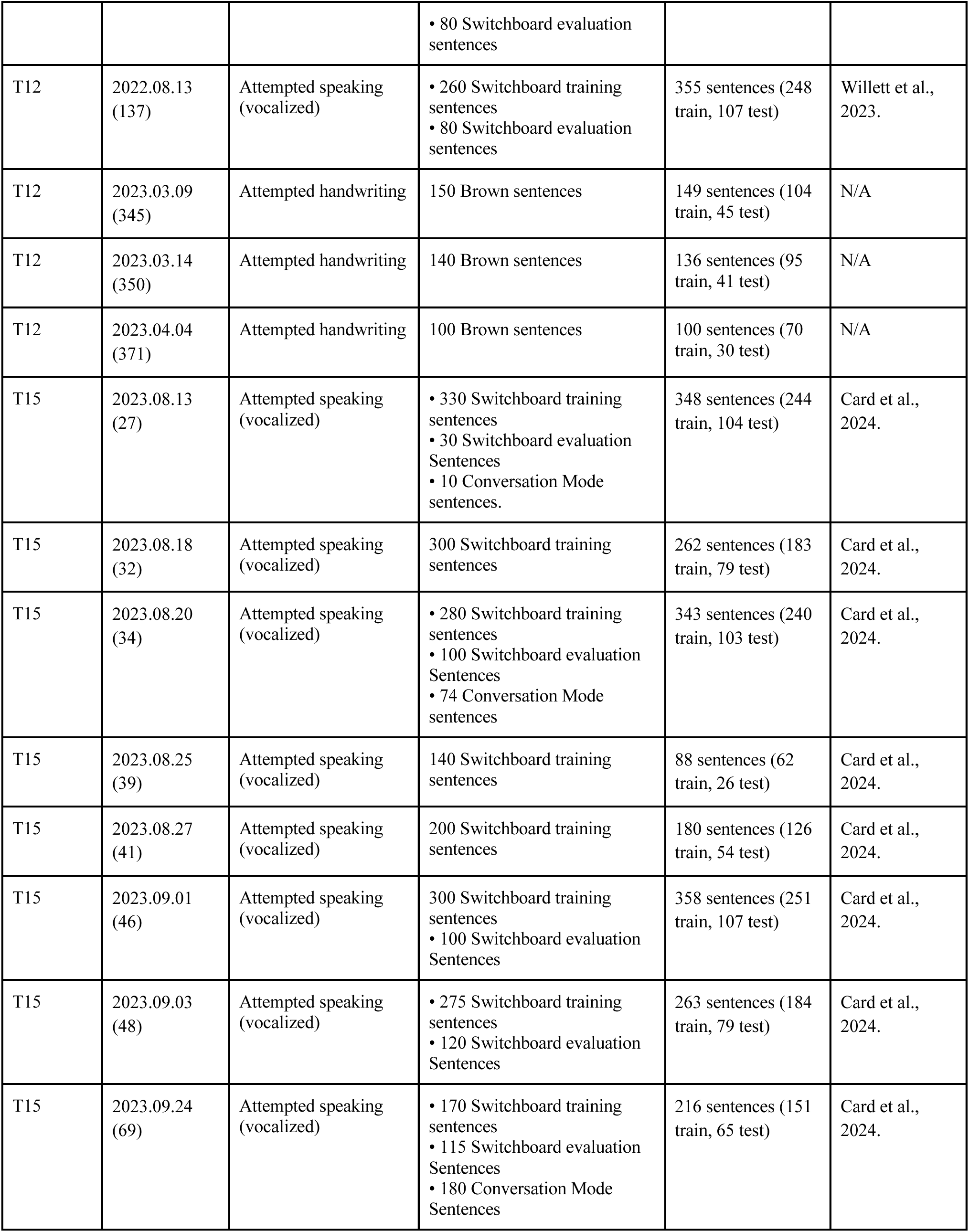
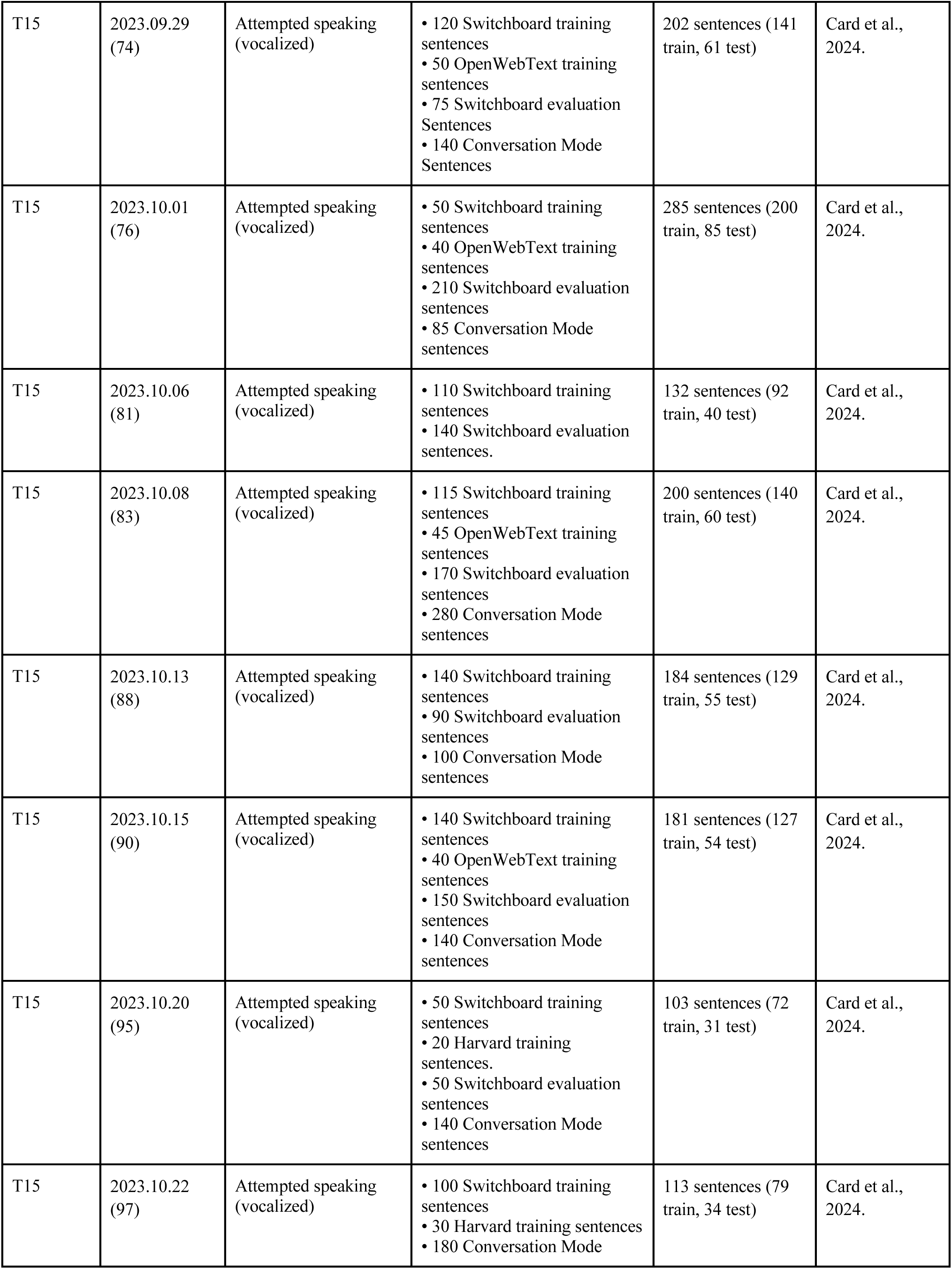
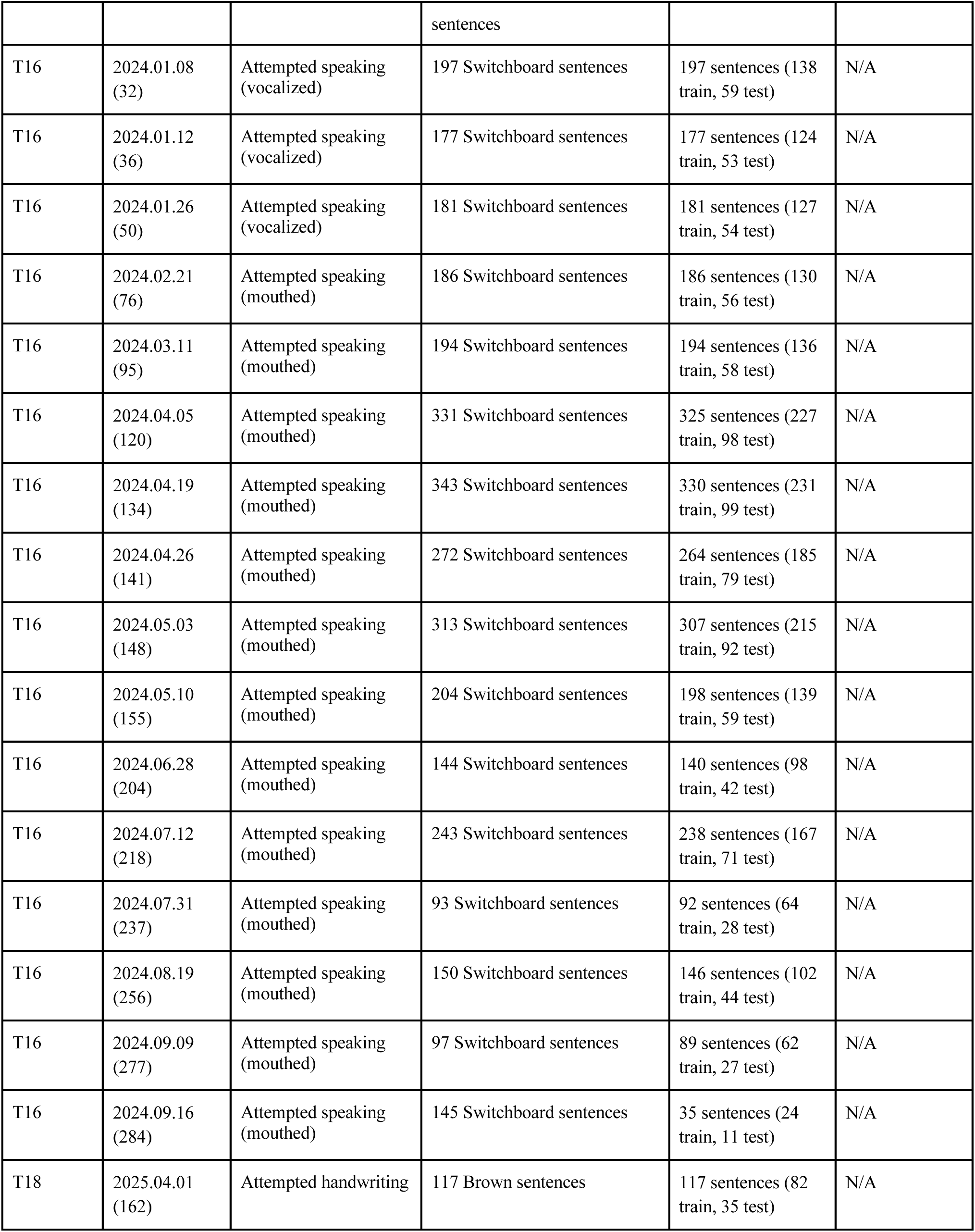
List of all data collection sessions included in this study.

### Data collection rig

Digital signal processing and feature extraction were performed on a dedicated computer. For all T5 sessions, and for T12 sessions conducted prior to December 2024, data was processed using Simulink Real-Time, and task software was implemented with the MATLAB Psychophysics Toolbox^35^. An additional Windows computer controlled the starting and stopping of tasks and interfaced with the Neuroplex-E system (Blackrock Microsystems). BRAND^28^, a framework for developing modular, Python-based neural data processing and task software, was used for all T12 sessions conducted from December 2024 onwards, as well as for all T15, T16, and T18 sessions.

### Instructed delay paradigm for both speech and handwriting

All data collection sessions employed an instructed delay paradigm, with each trial consisting of a “delay” period followed by a “go” period. During the delay period, a sentence was displayed on the screen above a red square, allowing the participant time to read it and prepare to either write or speak that sentence, depending on the task. After the delay period, the red square turned green to indicate the start of the go period. During this period, the sentence remained on the screen, while the participant attempted to write or speak the sentence (see Fig. 1a for task diagram). Once the participant had finished writing or speaking a prompted sentence, progression to the next trial in the experimental block could be triggered by: the participant pressing a button held in their lap (T12, T16), selecting a button using an eye tracker mounted on the screen or using gesture decoding (T15), turning their head to the right, which was detected by an OptiTrack system tracking the position of optical markers worn on their handband (T5), or by the researcher pressing a button on the participant’s behalf (T18). The speech session was typically composed of blocks of 40–50 trials, while the handwriting sessions consisted of 10-trial blocks. Between blocks, all participants were encouraged to rest as needed. All data collection sessions were performed at the participant’s place of residence.

During all speech sessions used in this study, participants T12 and T15 attempted to produce voiced speech, trying to move all of their articulators and modulating their larynx to pass sound, as one would typically do when attempting to speak. T16, on the other hand, primarily used “mouthed” or silent speech (as in Willett et al., 2023), producing no audible sound, but visibly attempting to move her lips, tongue and jaw as though mouthing to someone across the room.

In all handwriting sessions, participants were instructed to attempt to write the sentence letter by letter, with each letter superimposed on the previous one. Greater-than symbols (“>”) were used to signify spaces between words, while tildes (“∼”) were used to signify periods at the end of a sentence. Since T12 retains partial use of her limbs and communicates primarily through the use of an iPad tablet, she was able to write on a 15-inch LCD writing tablet (ERUW Shenzhen Lei Rui Technology Co., Ltd.) using a stylus during handwriting sessions (see Kunz et al., 2025 for details on the tablet setup). T5 and T18, both being tetraplegic, attempted to write with imagined pen and paper.

Each speech session began with either a speech-specific “diagnostic” block or a general “rest” block, which was used to calculate the threshold values and filters for linear regression referencing (LRR) to be applied throughout the session for T12 and T16. For T15, filter and LRR parameters were further recalculated after each block. Each handwriting session similarly began with a “rest” block that was used to calculate the threshold values and filters for linear regression referencing (LRR) to be applied throughout the session for T5, T12 and T18.

The majority of the speech sentences were collected in “open-loop” blocks as training data, with no real-time decoder feedback. However, some sentences—particularly toward the end of experimental sessions—were collected in “closed loop,” with real-time speech decoder output during each trial. All of the handwriting sentences used in this study were collected in “open loop” blocks.

In addition to data collection sessions that were conducted expressly for the investigation of cross-brain transfer, this work leveraged previously collected speech and handwriting data that have been made publicly available with the publication of prior work^3,5,36^. For a summary of all data collection sessions, see Table I.

### Sentence selection

For the speaking task, sentences were sourced from the Switchboard corpus of telephone conversation transcripts between speakers of American English^37^; see Willett et al., 2023 for details of the manual sentence screening process. Additional sentences were drawn from the OpenWebText2 corpus^38^ and the Harvard Sentences^39^. See Table I for a detailed breakdown of sentences collected from each corpus. For the handwriting task, sentences were drawn from the Brown Corpus, a collection of American English prose published in the United States in 1961^40^.

### Decoder training data

#### Neural feature pre-processing

Threshold crossing rates and spike band power features were binned into 20 ms time steps, z-scored (mean-subtracted and divided by the standard deviation), causally smoothed by convolving with a Gaussian kernel (sd = 40ms) that was delayed by 160ms, and concatenated into a 512 x 1 vector for speech data, and a 768 x 1 vector for handwriting data.

#### Held-out target user evaluation days

For speech, we designated one held-out session each for all three speech participants to use as a simulated day of initial BCI use (t12.2022.05.26, t15.2023.10.01, t16.2024.01.12). For each of the three possible target users, data from the other two source users were used to train a “base” decoder, and the held-out session from the target user was then used to fine-tune that decoder. For handwriting, only participant T5 had a large library of handwriting data. For participant T18, there was only one handwriting session (t18.2025.04.01), while for participant T12 there were three, one of which was designated as the held-out handwriting session (t12.2023.03.14). Therefore, we tested only participants T12 and T18 as target users, each paired with the single source user T5.

#### Train and test splits

For each session, the pre-processed trials were randomly shuffled across all blocks, and then split into 70% train and 30% test sets on which the decoder could be trained and evaluated. Additionally, since some sentence prompts had been reused across participants in prior work, we excluded these duplicates if they appeared in the held-out target day test set, in order to ensure that cross-brain transfer was evaluated on entirely held-out sentences. Finally, for the held-out target days, we created progressively larger training set variants, ranging from 10% to 100% of the training set, in 10% increments.

#### Zero-padding to match feature length across participants

Since participants had different numbers of microelectrode array channels placed in brain regions relevant to speech or handwriting decoding, we zero-padded our features to a fixed size based on the participant with the largest number of electrodes. For speech sessions, features were padded to 512, matching the neural feature count of participant T15. For handwriting sessions, features were padded to 768, matching the neural feature count of participant T18.

#### Permuted data

We also created randomly permuted variants of all non-held-out sessions. For each participant, a single, random ordering of channels was generated, and then applied to each of that participant’s non-held-out sessions prior to generation of train and test set splits.

### Decoder performance metrics

To evaluate the output of the decoder, we compared the most probable phoneme/character decoded at each time step (with duplicates removed) to the ground truth phoneme/character sequence of the sentence prompt (greedy decoding).

#### Speech

We evaluate phoneme error rate, defined as the edit distance between the decoded sequence of phonemes and the phoneme transcription of the sentence prompt (i.e., the number of insertions, deletions or substitutions required to make the sequence of phonemes match exactly).

#### Handwriting

Character error rate was defined as the edit distance between the decoded sentence and the sentence prompt (i.e., the number of insertions, deletions or substitutions required to make the strings of characters match exactly).

Note that the reported error rates are the combined result of many independent sentences used for decoder evaluation. To combine data across multiple sentences, we summed the number of errors across all sentences, and divided this by the total number of phonemes/characters across all sentences (as opposed to computing error rates first for each sentence and then averaging those error rates). This helps prevent very short sentences from overly influencing the result. Lastly, we generated 95% confidence intervals for all reported phoneme and character error rates with error rates from multiple decoder seeds.

### RNN decoder

#### Architecture & CTC loss

The decoding and training methods in this study are the same as those reported previously^3,5^. To transform neural activity evoked by either speech or handwriting into a time series of phoneme or character probabilities, we used a 5-layer, stacked gated recurrent unit RNN as described in Willett et al. 2023. Prior work found that fitting unique, trainable input layers for each dataset significantly improved both handwriting^1^ and speech^3^ decoding performance across days for a single participant, combating changes in neural features over time. Thus, the RNN also retained this separate layer for each data collection session: an affine transformation applied to all of the neural feature vectors from that particular session, followed by a nonlinear activation function:

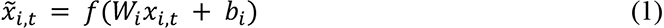

Here, *x_i,t_* is the input vector at time step *t* of session *i* and *x̄_i,t_* is the transformed input. *W_i_* ∈ ℝ*^p×p^* is a weight matrix and *b_i_* ∈ ℝ*^p^* is a bias vector for session *i*, with the number of input features, *p*, equal to 512 for speech data and 768 for handwriting data. The nonlinear activation function, *f* is then applied element-wise to the resultant vector, and here we use the softsign function *f*(*x*)/(|*x*|) + *1*. The weights and biases for each unique session were optimized simultaneously along with all other, shared RNN parameters. During training, dropout was applied both prior to and after the softsign.

Though the standard decoder architecture used throughout this work included session-specific input layers, we also used an ablated variant in which the unique layers were replaced by a single, shared input layer applied to the entire dataset, across all participants and session days.

Since all of the participants in the speech data collection sessions are unable to produce intelligible speech, and because two out of three of the participants in the handwriting data collection sessions are unable to physically write, we had to train the decoders without any ground truth timing labels. To achieve this, we used a connectionist temporal classification (CTC) loss^41^. We also added two types of artificial noise to help regularize the decoder: white noise was added directly to the input neural feature vectors at each time step in order to improve generalization, and artificial constant offsets were added to the means of the neural features, to make the decoder more robust to drifts in baseline firing rates that are known to accumulate over time and degrade decoding performance. The decoder ran at a 4-bin frequency (with 20 ms bins), outputting a phoneme or character probability vector for every 80 ms of neural activity. For more details on decoder implementation, see the supplemental methods sections of Willett et al. 2023 and Card et al., 2024. Certain hyperparameters differed between the speech and handwriting decoders; see Table II for a list of RNN hyperparameters used in the different experiments in this study.

**Table II:** Hyperparameters for all decoders included in this study. RNN hyperparameter spreadsheet (link)

#### Training data distributions

When training base decoders from scratch on data from one (handwriting) or two (speech) source users, we sampled from each day with uniform probability for all days. However, when base decoders were then fine-tuned on the target user’s held-out data, 30% of training was equally distributed across the source participants’ session days, while 70% of training was equally distributed across all target user days. We included the source user data during fine-tuning due to our prior work demonstrating the utility of a replay buffer^36^. Base decoders were evaluated on a uniform distribution of all test sets, while fine-tuned decoders were evaluated on only the test set of the target user’s test day.

## Acknowledgements

We thank participants T5, T12, T15, T16, and T18 and their care partners for their generous time and contributions to this research. We also appreciate the administrative support from B. Davis, K. Tsou, S. Kosasih, M. Massood, B. Travers, and D. Rosler, as well as Steve Mernoff for clinical site oversight.

This work was supported by an ALS Pilot Clinical Trial Award (AL220043) from the Office of the Assistant Secretary of Defense for Health Affairs; a New Innovator Award (NIH 1DP2DC021055) from the National Institutes of Health Office of the Director and managed by the National Institute on Deafness and Other Communication Disorders; NIH-NINDS U01NS123101; the NSF Graduate Research Fellowship (DGE-1656518) and the Bruce and Elizabeth Dunlevie Stanford Bio-X Interdisciplinary Graduate Fellowship (A.D.L.); NIH F32HD112173 (S.R.N.-T.); the NSF CAREER Award (2440859) and awards from the Sloan, Simons, and McKnight Foundations (S.W.L.); Larry and Pamela Garlick and the Stanford Wu Tsai Neuroscience Institute (J.M.H.); the Searle Scholars Program and has a Career Award at the Scientific Interface from the Burroughs Wellcome Fund (S.D.S); NIH-NINDS/OD DP2NS127291 and the Simons Foundation as part of the Simons-Emory International Consortium on Motor Control (C.P.).

The content is solely the responsibility of the authors and does not necessarily represent the official views of the National Institutes of Health, or the Department of Veterans Affairs, or the United States Government. CAUTION: Investigational Device. Limited by Federal Law to Investigational Use.

## Author Contributions

A.D.L. and F.R.W. conceived the study and led the interpretation of all experiments, and ADL wrote the manuscript and led the development and analysis of all experiments.

D.T.A. developed the data collection rig software at the Stanford University site, as well as built a tablet and stylus setup for T12 handwriting sessions.

N.S.C., M.W., B.G.J., and J.J.J. developed and executed the experiments at their respective sites.

F.B.K., C.I., P.H.B., S.R.N.-T., and B.E.L. were responsible for coordination of session scheduling, logistics and daily equipment setup/disconnection for participants T12, T15, T16, and T18 respectively.

D.R.D. compiled Human Connectome Project parcellation results to generate Figure 1.

L.R.H. is the sponsor-investigator of the multisite BrainGate2 pilot clinical trial.

J.M.H. planned and performed T5’s and T12’s array placement surgery and was responsible for all clinical trial-related activities at Stanford University.

D.M.B. planned and performed T15’s array placement surgery and was responsible for all clinical trial-related activities at University of California, Davis. S.D.S. and D.M.B. supervised and guided all research activities at UC Davis.

N.A.Y. planned and performed T16’s array placement surgery and was responsible for all clinical trial-related activities at Emory University. C.P. and N.A.Y. supervised and guided all research activities at Emory.

L.R.H., S.S.C. and Z.M.W. planned T18’s array placement surgery, and Z.M.W. performed T18’s array placement surgery. L.R.H. was responsible for all clinical trial related activity at VA Providence and Massachusetts General Hospital. L.R.H. and D.B.R. supervised and guided all research activity with T18.

The study was supervised and guided by S.L., J.M.H. and F.R.W. All authors reviewed and edited the manuscript.

